# Human Longevity is Influenced by Many Genetic Variants: Evidence from 75,000 UK Biobank Participants

**DOI:** 10.1101/038430

**Authors:** Luke C. Pilling, Janice L. Atkins, Kirsty Bowman, Samuel E. Jones, Jessica Tyrrell, Robin N. Beaumont, Katherine S. Ruth, Marcus A. Tuke, Hanieh Yaghootkar, Andrew R. Wood, Rachel M. Freathy, Anna Murray, Michael N. Weedon, Luting Xue, Kathryn Lunetta, Joanne M. Murabito, Lorna W. Harries, Jean-Marie Robine, Carol Brayne, George A. Kuchel, Luigi Ferrucci, Timothy M. Frayling, David Melzer

## Abstract

Variation in human lifespan is 20 to 30% heritable but few genetic variants have been identified. We undertook a Genome Wide Association Study (GWAS) using age at death of parents of middle-aged UK Biobank participants of European decent (n=75,244 with father’s and/or mother’s data). Genetic risk scores for 19 phenotypes (n=777 proven variants) were also tested.

Genotyped variants (n=845,997) explained 10.2% (SD=1.3%) of combined parental longevity. In GWAS, a locus in the nicotine receptor *CHRNA3* – previously associated with increased smoking and lung cancer - was associated with paternal age at death, with each protective allele (rs1051730[G]) being associated with 0.03 years later age at father’s death (p=3x10^-8^). Offspring of longer lived parents had more protective alleles (lower genetic risk scores) for coronary artery disease, systolic blood pressure, body mass index, cholesterol and triglyceride levels, type-1 diabetes, inflammatory bowel disease and Alzheimer’s disease. In candidate gene analyses, variants in the *TOMM40/APOE* locus were associated with longevity (including rs429358, p=3x10^-5^), but *FOXO* variants were not associated.

These results support a multiple protective factors model for achieving longer lifespans in humans, with a prominent role for cardiovascular-related pathways. Several of these genetically influenced risks, including blood pressure and tobacco exposure, are potentially modifiable.

## Introduction

Over the last several decades, genetic studies conducted *in-vitro* and using model organisms have produced a stream of exciting findings linking specific pathways to major effects on aging [1], including, for example, gene knockouts in nutrient sensing pathways that cause dramatic life extension in *C. elegans.* However, humans outlive laboratory models many times over and humans suffer from several conditions that don’t normally affect laboratory animals. Moreover, beyond obvious differences in rate of aging, body size and higher brain functions, genomic responses in mice poorly mimic human inflammatory responses [2]. It is therefore uncertain to what extent the laboratory findings and the ‘single pathway major effect’ model is relevant to human aging and longevity [3]. Additionally, in humans, health behaviours plus social and economic factors play a major role in explaining differences in survival, with smoking being a substantial negative factor [4].

In addition to environmental exposures, genetic variation is important: for example, studies conducted on heterozygotic and homozygotic twins consistently report heritability of longer human lifespans of 20-30% [5,6], with negligible heritability before age 60 years but increasing estimates at advanced ages [7]; the low early heritability is hypothesized to be due to more “accidental” or exposure-related deaths at younger ages. Well-powered genome wide association studies (GWAS) provide robust evidence on which common single nucleotide polymorphisms (SNP) are associated with traits including longevity, but thus far sample sizes have been relatively modest. In the recent meta-analysis of data from 6,036 participants of European descent who survived beyond 90 years versus 3,757 younger controls (aged 55 to 80 years) [8], no genome wide significant variants were identified, but in candidate analyses the *APOE* locus was consistently associated across the participating cohorts. In this GWAS data evidence was found to support the *FOXO3* gene variants linked to longevity in candidate studies in Asian and other populations (best performing variant in discovery analysis rs10457180, uncorrected p-value=0.012 for *a priori* SNP rs2802292). In smaller previous studies, the frequency of proven disease risk increasing SNP alleles appeared no different in long lived individuals, which was interpreted as showing that longevity is achieved independent of disease risk alleles [9,10].

Identifying genetic variants associated with longevity using unbiased methods is challenging. As described above, a common ‘direct’ approach has been to compare older study volunteers with younger participants from the next generation, but this approach may be biased by the many changes in exposures (e.g. early infectious diseases, changing medical treatment) and increasing life expectancy across the generations. Ideal comparisons might be of exceptionally old individuals with those from a representative sample of their own generation who died at younger ages, but obtaining DNA samples from controls who died decades earlier is challenging. An ‘indirect’ approach to the ideal design, based on the assumption that longevity is a genetic transmissible trait, is to access DNA from offspring and test for variations associated with the longevity of their parents. The middle-aged offspring of long-lived parents have less cardiovascular disease, cancer, diabetes, and all-cause mortality compared to offspring whose parents died at younger ages [11], consistent with the inheritance of longevity associated genetic variants. This better health status in offspring showed a linear association with advancing parental age, with a slightly stronger association with mother’s compared to father’s age at death. Given that offspring inherit their DNA from two parents who might have died at very different ages, associations with longevity in offspring are diluted and samples 3–4 times larger than the direct younger vs older approach are needed. In fact, Tan et al [12] estimate that 1,500 participants of at least one long-lived parent would be needed to achieve >90% power to detect less common alleles (5% frequency) with effects of 0.85 (in binary analysis of offspring of 1 long-lived parent vs. controls) with 95% confidence.

In the current analysis we aimed to identify common genetic variants (prevalence ≥1%) associated with longer parental lifespan. To achieve the sample sizes required, we utilized data from UK Biobank. We first performed genome-wide association studies (GWAS) and then used genetic risk scores (GRS) of known variants to test the hypothesis that offspring of longer-lived parents have lower genetic-risk of common risk traits and diseases. In these analyses we have included middle aged participants only (age 55 to 70), as typical intergenerational age gaps mean that the parents of the younger UK Biobank participants were definitely or likely to be too young to have achieved longer survival, at the time of the UK Biobank baseline interview.

## Methods

Between 2006 and 2010, 503,325 volunteers (aged 45 to 69 years old) were recruited from across the United Kingdom to the UK Biobank study [13]. Of these, 75,244 participants met the inclusion criteria for at least one analysis: participants aged between 55-70 years with complete genetics data and date of death data for either parent (additional participants were included if they had an alive parent that met the “long-lived” criteria for binary analyses). We chose this 55 to 70 age-range because usual intergenerational age gaps mean that very few participants below age 55 at baseline could have parents old enough to be longer lived, thus including younger ages would enrich and potentially bias our study toward premature and early deaths. Participants were excluded if they reported themselves as adopted, or if either parent died prematurely (fathers <46 years or mothers <57 years – see methods below), or if their parent was still alive but not yet long-lived. Several longevity phenotypes are defined below based on the age at death of the participant’s parents.

### Parental age at death and longevity phenotypes

Participants were asked the age at which their parents had died (or their current age if still alive). Analyses were performed separately on mother’s age at death and father’s age at death, and also on a combined phenotype. To reduce the effect of higher ages at death of mothers (compared to the fathers) we first z-transformed the mothers and fathers age at deaths before combining the z-scores into a single summed phenotype. Offspring of parents who died prematurely were excluded because the cause of death of the participant’s parents was not asked, so we could not exclude accidental deaths explicitly. To determine the premature age at death cut-offs we used previously described methods to define the normal range of age at death for mothers and fathers separately, and excluded participants below these values (methods described here [11], description of method applied to UK Biobank here (Atkins et al. in submission 2016)). The analysis identified the following cut-points for short, intermediate, and long-lived parents ages at death: for mothers (57 to 72 years, 73 to 92 years, and ≥93 years, respectively) and for fathers (46 to 65 years, 66 to 89 years, and ≥90 years, respectively). Parents below the “short” definition are excluded as premature deaths.

We also defined a binary trait for an “extreme longevity” phenotype by determining the top 1% of age at death for mothers (≥98 years) and fathers (≥95 years) separately. We were able to increase the number of “long lived” parents in all the binary phenotypes by including participants who had responded to the question regarding the age of their parent if still alive. Participants with one long-lived and one short-lived parent were excluded from this analysis. Finally, we defined two further binary traits for sensitivity analyses testing the consistency of results using a non-linear definition: offspring of at least one long-lived parent vs. offspring of two intermediate-lived parents, and offspring of at least one long-lived parent vs. offspring of at least one short-lived parent (very low numbers of participants with either “both long” or “both short” meant we used the aforementioned “at least one…” definitions). **Supplementary Figure 1** contains a flow-chart showing how the phenotypes are derived and the numbers included in each analysis.

### UK Biobank genetics data

We used genetic data available (autumn 2015) from 120,286 participants identified as ‘white British’ through self-report and verified through principal components analysis based on genotypes. Kinship coefficients were estimated and related individuals (3rd degree or higher) were removed to provide the maximal unrelated set of individuals. The central UK Biobank analysis team performed these analyses. Details of principal component analyses and kinship analyses can be found in the official UK Biobank genotyping document (http://biobank.ctsu.ox.ac.uk/crystal/docs/genotyping_qc.pdf; accessed 1st December 2015).

We used imputed genotypes available from the UK Biobank for association analyses. Briefly, phasing of individuals was carried out using SHAPEIT-2. Imputation was performed using IMPUTE2 and a combined 1000 Genomes / UK10K reference panel. Full details can be found in the official UK Biobank imputation document (http://biobank.ctsu.ox.ac.uk/crystal/docs/impute_ukb_v1.pdf; accessed 1st December 2015). After filtering for variants with MAF ≥1%, missingness <1.5%, imputation quality >0.4 and with Hardy-Weinberg equilibrium (HWE) P>1x10^-6^ within the white British participants, 9,658,292 imputed autosomal variants were eligible for the analyses.

We also utilized data directly from the microarrays for variants on the X (n=19,381) and Y (n=284) chromosomes, and on the mitochondrial genome (n=135), which were unavailable in the imputed dataset.

### Within “white British” principal components

We selected 95,535 independent SNPs (pairwise r2<0.1) directly genotyped with a minor allele frequency (MAF) ≥ 2.5% and missingness <1.5% across all UK Biobank participants with genetic data available at the time of this study (n=152,732), and with HWE P>1x10-6 within the white British participants. Principal components were subsequently generated using FlashPCA and the first five adjusted for in all analyses [14].

### Power Calculations

Quanto software version 1.2.4 was utilized for power calculations with parameters: continuous, additive model, mean/SD of standardized outcome=0/1 [15].

### Genome Wide Association Study

We used BOLT-LMM to model the associations between imputed variants (dosages) and each phenotype [16], which uses a linear mixed effects model approach. We looked at the results for variants with imputation quality >0.4, HWE p-values >1x10^-6^ and minor allele frequencies >0.1% in the white/British subset used for all analyses. For variants on the X, Y and mitochondrial chromosomes only in the directly-genotyped data we used Plink (v1.9) [17] in linear (additive) or binary (fisher) models, as appropriate, adjusted for the same covariates as above including the first 5 principal components from FlashPCA.

### Independent testing of GWAS results

We obtained estimates of our genome wide significant results for fathers age at death GWAS in the Framingham Heart Study (FHS) generation 2 [18], with data on n=2033 offspring available. The inclusion criteria and model specification were the same as described for Biobank, with the exception that family structure was taken into account (using R package ‘pedigreem’).

### Genetic Heritability Estimation

To estimate the variance in mean parental age at death (and age at death of mothers/fathers considered separately) explained by common genetic variants we utilized the BOLT-REML package [19]. This package uses REstricted Maximum Likelihood (REML) methods for variance component estimation, in this case on the genetics data. We used SNPs that met the following criteria in the 120,286 individuals; variants were excluded if not in Hardy-Weinberg Equilibrium (HWE) (P<1x10-6), or had a minor allele frequency <1%, or an overall missing rate >1.5% in any individual batch, or were on the Y chromosome. This resulted in 457,643 directly genotyped variants for inclusion. We also performed an additional sensitivity analysis relaxing the exclusion criteria to include all 845,997 variants. BOLT-REML determines the “heritability” of phenotypes based on the variance components of the genetics data provided. Prior to analyses, phenotypes are adjusted (by taking the residuals from a linear regression analysis) for confounding factors (age, sex, array type, assessment center, principal components 1-5).

### Genetic Risk Score (GRS) creation

For each trait we identified the most recent GWAS meta-analysis and downloaded the results tables. We selected the SNPs identified at genome wide significance (p<5x10^-8^) and extracted the corresponding genotype information from the imputed data in the UK Biobank data. For all SNPs included in a GRS we checked for high imputation quality (>0.9), no significant deviation from HWE (p>1x10^-6^) and low missingness (<5% sample missing). We used the imputed genotype (dosage) information and effect size (from the meta-analysis) with Plink (v1.9) function ‘scores’ to generate the weighted GRS for each trait in each participant [17]. The Plink function uses genotype coding for each participant as 0, 1 or 2 trait-increasing alleles, which is multiplied by the effect size (coefficient or odds ratio) from the published study. The resulting “weighted allele score” is summed for all variants associated with a particular trait. See the accompanying **Supplementary Methods** document for the definition of each genetic risk score, and **Supplementary Table 1** for the variants and effect sizes used to create the scores.

### Statistical Analysis of Genetic Risk Scores

Generalized linear regression models were used throughout (R v3.2.0), with adjustment for age, sex, array type (‘axiom’ or ‘bileve’), assessment center (22 possible categories), and the first 5 genetic principal components. The regression linker functions ‘logit’ or ‘Gaussian’ were used respectively for logistic and linear outcomes. Forest plots comparing the associations between different GRS’ and each outcome were generated using the R package ‘rmeta’ (v2.16). For these analyses the GRS was first z-transformed so that the statistics reported could be compared between each GRS on the forest plot. In the rest of the manuscript, we refer to the unstandardized, “per weighted allele” effects. In all respects except for the z-transformation the models were identical. Benjamini-Hochberg p-values are calculated to correct for potential false-positive associations (19 GRS against 4 primary longevity phenotypes).

## Results

Our sample (Table 1) included ‘white’ British UK Biobank participants aged 55-70 years old (n=75,244 with data on fathers survival, mothers survival or both; **Table 1).** There were relatively few current smokers (8.3% overall), but smoking rates were higher in those with short lived parents. The mean age at death of fathers was 72.9 (range 46 to 105 yrs) and mothers 78.5 years old (range 57 to 107 yrs). Nearly half the participants (48.5%) were men. Three continuous phenotypes were utilized throughout this analysis; participant’s father’s age at death (n=63,775), mother’s age at death (n=52,776) and combined (normalized) mothers and fathers ages at death (n=45,627 with age at death data for both parents). We had 90.4% power to detect an allele of 5% minor allele frequency in the sample of 45,627 accounting for 0.1% of the variance in the phenotype after multiple-testing correction (alpha=5x10^-8^).

**Table 1:** Summary statistics for the UK Biobank participants eligible for at least one analysis

| Phenotype | *All participants* |  |  | *Longer-lived parents ‡* |  |  | *Medium-lived parents ‡* |  |  | *Short-lived parents ‡* |  |  | p-value * |
| --- | --- | --- | --- | --- | --- | --- | --- | --- | --- | --- | --- | --- | --- |
|  | N | Mean | SD | n | Mean | SD | n | Mean | SD | n | Mean | SD |  |
| Age at recruitment (yrs) | 75,244 | 62.077 | 4.060 | 8,655 | 63.529 | 3.659 | 21,299 | 62.862 | 3.883 | 25,771 | 62.096 | 4.039 | 3.6 x10<sup>-209</sup> |
| Body mass index (BMI) | 75,038 | 27.654 | 4.687 | 8,633 | 26.970 | 4.358 | 21,248 | 27.537 | 4.588 | 25,683 | 28.040 | 4.812 | 2.1 x10<sup>-80</sup> |
| Fathers age at death (yrs) | 63,775 | 72.925 | 11.095 | 7,171 | 81.844 | 12.456 | 21,299 | 76.973 | 6.143 | 23,984 | 64.396 | 10.209 | n/a |
| Mothers age at death (yrs) | 52,776 | 78.473 | 9.489 | 6,043 | 87.215 | 9.789 | 21,299 | 82.130 | 5.269 | 21,422 | 71.808 | 9.009 | n/a |
| Combined normalized parent age death ° | 45,627 | 0.003 | 1.526 | 4,052 | 2.318 | 0.697 | 21,299 | 0.736 | 0.818 | 19,635 | -1.336 | 0.978 | n/a |
| Sex (% males) | 75,244 | 48.49% | n/a | 8,655 | 49.10% | n/a | 21,299 | 48.72% | n/a | 25,771 | 48.79% | n/a | 8.3 x10<sup>-1</sup> |
| Smoking status (% current) | 74,990 | 8.34% | n/a | 8,637 | 6.24% | n/a | 21,237 | 7.78% | n/a | 25,671 | 8.98% | n/a | 7.5 x10<sup>-16</sup> |

‡ "Long-lived"=offspring of at least one long-lived parent, "medium-lived"=two intermediate-lived parents, "short-lived"= at least one short-lived parent
\* p-value from Kruskal-Wallis non-parametric analysis of variance test for significant difference in distribution of phenotype between the 3 longevity categories
° Z-scored mother's and father's ages at death, summed

In addition we created a binary “extreme longevity” phenotype based on the top 1% of the age at death distribution for mothers and fathers (≥98 years and ≥95 years, respectively). Of 45,627 participants with age at death data for both parents 907 had at least one parent who died within these ranges (1.99%). An additional 432 participants had at least one parent still alive who met the criteria; therefore 1,339 participants had a least one parent (alive or dead) who lived to an “extreme age” (3.2% of 42,273 participants; those with ‘discordant’ parents – one long-lived and one short-lived – are excluded). See **Supplementary Figure 1** for a flowchart of the participants included in each of the four primary analyses.

### Genome Wide Association Study: smoking-related variants are associated with father’s age at death

Of the 9,658,292 genetic variants included none were significantly associated (p<5x10^-8^) with “parent’s age at death, combined” however one locus (36 variants in strong Linkage disequilibrium) on chromosome 15 was associated with “father’s age at death,” and 1 variant (on chromosome 22) with “mother’s age at death” (see **Supplementary Table 2** for the top 1,000 results from each GWAS; see **Supplementary Figures 2-5** for Manhattan and QQ plots). We focused on the locus on chromosome 15 associated with father’s age at death, because the variant associated with mother’s age at death was of low frequency (3%) and not in a typical “peak” expected of robust results (see **Supplementary Figures 6-7** for LocusZoom plots [20]). rs1051730 is in this loci (beta between G allele and father’s age at death=0.0269, se=0.0049, p=3x10^-8^); the A allele of rs1051730 in the nicotinic acetylcholine receptor alpha 3 subunit *CHRNA3* gene has been linked to smoking fewer cigarettes and lower risks of lung cancer, although this variant does not influence the chances of starting to smoke [21].

The association between rs1051730 and mother’s age at death (Beta=0.017, p=1.6x10-3) did not reach genome wide significance in our UK Biobank participants. We tested the association of rs1051730 with father’s age at death in the Framingham Heart Study (FHS) generation 2, where 2033 participants were available (mean age at death of the fathers=77.4 years, SD=11.5, range=47.2 to 102.8). The association was non-significant, but was directionally consistent (per C allele coefficient= 0.008, p=0.98). However, the effects size of this SNP is modest and the power to detect the association in the FHS was only 1% (Quanto parameters: coefficient=0.0269, n=2033, MAF=0.17, alpha=0.05).

In the GWAS against the binary “extreme age” phenotype (at least one parent very long-lived) two variants (rs528161076 and rs75824829, on chromosome 7 and 9 respectively (see **Supplementary Figures 8-9** for Locus Zoom plots) were found to be significant (p<5x10^-8^). rs75824829 may be anomalous as it is not in a “peak” of associations expected of robust results **(Supplementary Figure 4; Supplementary Figure 9),** however rs528161076 is in a distinct peak of variants in an intron of *AP5Z1 (adaptor related protein complex 5, zeta 1 subunit,* thought to be involved in homologous recombination DNA doublestrand break repair). The association between rs528161076 and extreme longevity should still be interpreted with caution, as the variant is not associated with the continuous age at death variables (combined parent’s age at death p=0.65, father’s p=0.47 and mother’s p=0.96).

In all four GWAS performed, no variants on the X or Y chromosomes were associated with the phenotypes (p>1x10^-5^). For mitochondrial variants, the smallest p-values for mother’s age at death, was 9x10^-3^.

### Heritability of parental longevity explained by genotyped common variants

We determined the variance in parental age at death explained by all the common genetic variation (minor allele prevalence >1%) robustly genotyped directly on the UK Biobank arrays (n=457,643; see methods). For combined parents age at death, the variance attributed to the measured genotypes was 8.47% (SD=1.06%); for mothers age at death 4.85% (SD=1.01%) and fathers age at death 5.35% (SD=1.04%). In a sensitivity analysis including all directly genotyped variants irrespective of minor allele frequency and missingness (n=845,997) the variance attributed for the combined age at death phenotypes was 10.24% (SD=1.26%); for mothers age at death 6.08% (SD=1.21%) and fathers age at death 5.79% (SD=1.23%).

### Genetic Risk Score associations

For each genetic risk score (GRS) we first verified the association with the best-fit phenotype corresponding to the reported association: for example the coronary artery disease (CAD) GRS was tested against prevalent coronary heart disease in the offspring (CHD= myocardial infarction or angina) **(Table 2).** All GRS were associated with their available phenotypes in UK Biobank in the expected direction, for example CAD GRS and CHD (per weighted allele increased OR=1.12, 95% CIs: 1.10 to 1.13, p=1x10^-64^). The lipid GRS could not be tested directly, as serum lipid concentrations are not yet available, however all three separate risk scores are significantly associated with CHD. Although there were only 28 participants with dementia, a significant association was seen with the Alzheimer’s disease GRS (per weighted allele OR=1.52, 95% CIs: 1.17 to 1.98, p=2x10^-3^). Telomere length was not assessed in the Biobank participants so validation was not possible.

**Table 2:** Genetic Risk Score associations with corresponding phenotypes in 75,244 participants included for at least one analysis

| Model | GRS (n SNPs) | Phenotype (n cases) | OR (95% CIs) | p-value |
| --- | --- | --- | --- | --- |
| *Logistic regression models* |  |  |  |  |
|  | Coronary Artery Disease, CAD (53) | Prevalent CHD (5067) | 1.12 (1.10 to 1.13) | 2.3 x10-63 |
|  | CAD, without LDL, HDL and TG SNPs (25) | Prevalent CHD (5067) | 1.13 (1.11 to 1.15) | 6.1 x10-36 |
|  | LDL cholesterol (49) | Prevalent CHD (5067) | 2.97 (2.40 to 3.68) | 1.3 x10-23 |
|  | HDL cholesterol (67) | Prevalent CHD (5067) | 0.62 (0.50 to 0.78) | 3.1 x10-05 |
|  | Triglycerides, TG (37) | Prevalent CHD (5067) | 1.88 (1.43 to 2.46) | 5.3 x10-06 |
|  | Stroke (4) | Prevalent stroke or TIA (1790) | 2.04 (1.03 to 4.06) | 4.0 x10-02 |
|  | Type-1 Diabetes (29) | Type-1 diabetes diagnosis (55) | 2.35 (1.98 to 2.78) | 9.8 x10-23 |
|  | Type-2 Diabetes (55) | Type-2 diabetes diagnosis (4,052) | 5.29 (4.55 to 6.16) | 1.7 x10-103 |
|  | Alzheimer's Disease (8) | Dementia diagnosis (28) | 1.52 (1.17 to 1.98) | 1.9 x10-03 |
|  | Inflammatory Bowel Disease (156) | Inflammatory Bowel Disease (654) | 1.13 (1.10 to 1.16) | 5.3 x10-22 |
|  | Crohn's Disease (139) | Prevalent Crohn's disease (250) | 1.16 (1.11 to 1.21) | 1.1 x10-12 |
|  | Ulcerative Colitis (87) | Prevalent ulcerative colitis (414) | 1.17 (1.13 to 1.22) | 4.4 x10-15 |
|  | Prostate Cancer (85) * | Prevalent prostate cancer (863) | 1.18 (1.16 to 1.21) | 2.5 x10-52 |
|  | Breast Cancer (65) * | Prevalent breast cancer (2036) | 1.12 (1.11 to 1.14) | 2.3 x10-43 |
|  | Telomere Length (7) | n/a |  |  |
| *Linear regression models* |  |  | **Coefficient (95% CIs)** | **p-value** |
|  | BMI (69) | Body mass index | 7.07 (6.62 to 7.51) | 2.8 x10-209 |
|  | Systolic Blood Pressure (26) | Systolic blood pressure | 1.31 (1.14 to 1.47) | 4.3 x10-57 |
|  | Age at menopause (52) * | Age at menopause | -1.79 (-1.92 to -1.65) | 2.4 x10-154 |
|  | Forced Vital Capacity (6) | Forced Vital Capacity | 0.001 (0.001 to 0.002) | 7.0 x10-09 |

GRS = genetic risk score. OR = per weighted allele odds ratio. CIs = confidence intervals.
\* Analysis performed in male/female participants only, as determined by the phenotype

We next tested associations between each GRS and parental age at death (combined mothers and fathers) in linear regression models plus ‘extreme longevity’ (at least 1 parent lived to the top 1% of the age at death distribution in UK Biobank) **(Figure 1).** These analysis included 76 statistical tests (19 GRS against 4 primary longevity phenotypes) and therefore we have ‘starred’ associations passing Benjamini-Hochberg correction only, although each test has a strong and distinct prior hypothesis (see full results in **Supplementary Table 3).** We observed associations in the expected directions (i.e. lower disease risk score or higher numbers of relatively protective alleles associated with older ages at death) between the combined parental age at death continuous trait and eight GRS’; for CAD (unadjusted p=3.5x10^-18^), LDL cholesterol (p=1.8x10^-7^), CAD without lipid-associated alleles (p=6.56x10^-9^), triglycerides (p=1.0x10^-2^), systolic blood pressure (SBP, p=1.2x10^-4^), BMI (p=8.2x10^-4^), Inflammatory Bowel Disease (p=7.0x10^-3^), T1D (7.5x10^-3^), and Alzheimer’s disease (p=1.4x10^-2^). Crohn’s disease (p=2.3x10^-2^) and breast cancer (p=3.5x10^-2^) were associated at nominal significance (p<0.05) but not after multiple-testing correction.

**Figure 1.**
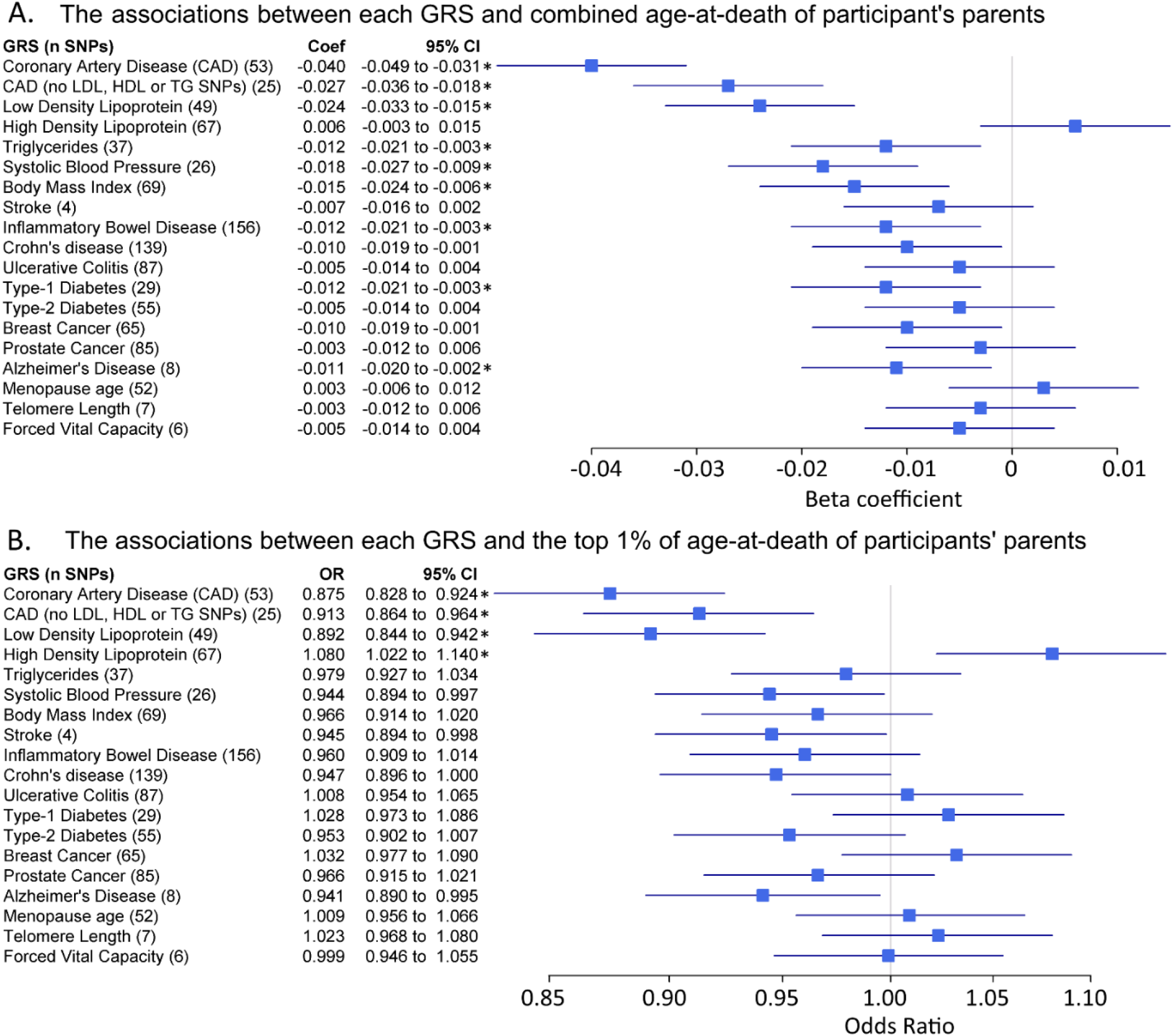
Forest plots show the relationship between the genetic risk scores (GRS, z-transformed) and 2 different parental longevity traits. Results from regression models (see methods). * indicates the association is significant after adjustment for multiple testing. A) Linear regression results for combined parent’s age at death. N=45,627. Coefficients (‘coef’) show the standard deviation (SD) difference in GRS per SD of parent’s age at death. B) Logistic regression results for the binary trait “participants of at least one parent reaching the top 1% of the age-at-death distribution vs. participants who parents did not”. Discordant participants (i.e. one long-lived and one short-lived) were excluded. n=42,273. n=1,339 participants of at least one long-lived parent (mother aged ≥98 years or father aged ≥95 years). Includes participants with alive parents reaching these limits. Odds Ratios (OR) show the likelihood of participants having at least 1 parent reaching the top 1% of age-at-death distribution per SD difference in GRS.

A number of GRS had similar trends for the extreme longevity association but become statistically non-significant in the binary analyses of offspring of long-lived parents, which has reduced statistical power (binary analysis vs. continuous). The HDL cholesterol GRS was associated with extreme longevity (p=5.7x10^-3^) although this association did not reach statistical significance in the linear analysis of continuous age at death.

We observed consistent effect directions, but reduced effect size and significance in separate analyses of mother’s age at death compared to father’s **(Supplementary Table 3);** with the exception of Alzheimer’s disease (AD) GRS which is only associated with mother’s age at death but not father’s (coefficient=-0.111, p=3.9x10^-4^; coefficient=-0.001, p=0.99, respectively) and Triglyceride level (TG) GRS was only associated with fathers not mothers age at death (father’s coefficient=-1.366, p=9x10^-4^; mother’s coefficient=-0.183, p=0.63, respectively). In linear regression models for AD and TG GRS there were no statistical interactions between mothers and fathers ages at death (p>0.05).

To illustrate combined effect sizes, we computed an estimate across the three GRS most robustly associated with parental longevity (unadjusted p<0.001): CAD, SBP and LDL. In total, 525 participants (of 45,627) were in the bottom quintile (20%) for all three GRS; these participants are the “lowest risk” group for these three cardiovascular traits. Correspondingly, 524 participants were categorized as having the “highest risk” (top 20% of genetic risk for all three traits). In adjusted logistic regression models, participants with the lowest genetic risk (highest number of protective alleles) had 3.25 times the likelihood of having at least 1 parent in the top 1% of ages at death, compared to those with the highest genetic risk (OR=3.25, 95%CI=1.3 to 8.1, p=0.012). Unadjusted prevalence: 19 participants (1.4% of 1,339) with at least one long-lived parent were in the bottom 20% for all three risk scores, where only 7 participants (0.5% of 1,339) were in the top 20% for all three risk scores.

### Candidate genetic variants associated with longevity

We have highlighted the GWAS results for a number of SNPs *a priori* selected from the literature (see **Supplementary Table 4** for the full details). We found significant (p<0.05) associations between 4 of the 5 *APOE/TOMM40* SNPs tested and parents age at death (rs2075650 p=1x10^-4^, rs429358 p=8x10^-7^, rs7412 p=0.02, rs4420638 p=3x10^-5^, but not rs405509 p=0.06). Variants rs2075650, rs429358 and rs4420638 are in linkage disequilibrium (R^2^ >0.4). Of 12 *FOXO3* variants assessed, none were significantly associated with a longevity phenotype (p>0.05). We observed modest associations with 2 of 7 variants at the *CDKN2A* locus (variants previously associated with coronary heart disease) with parent’s age at death (rs1333049 G allele beta=0.015, p=0.0015; rs4977574 A allele beta=0.014, p=0.0018); the *CDKN2A* variants associated with type-2 diabetes were not associated with parent’s longevity. Four variants were reported by Fortney to be associated with exceptional longevity; one is the *APOE* variant already discussed, and the other three are rs4977756 *(CDKN2B/ANRIL),* rs3184504 *(SH2B3/ATXN2)* and rs514659 *(ABO).* Only rs3184504 is associated with extreme longevity in this analysis (T allele beta=-0.0019, p=0.024), but rs4977756 was associated with continuous parent’s age at death (G allele beta=0.017, p=0.025). The final SNP is not associated with any longevity phenotypes in our analysis.

## Discussion

We report the largest analysis thus far of common genetic variants and human longevity, based on the indirect approach of testing variant associations in offspring. We identified a genome-wide significant variant in the smoking-related *CHRNA3* gene with father’s longevity. We also identified a variant associated with extreme longevity, in the *AP5Z1 gene* locus, a gene encoding a homologous recombination DNA double-strand break repair helicase protein [22], although this association with longevity appears less robust. In addition, we have shown that longevity is associated with having greater numbers of relatively protective alleles (i.e. lower genetic risk scores) for several cardiovascular traits including lipid levels, CAD (excluding variants associated with LDL and HDL), BMI, and blood pressure. Weaker associations were also found for risks scores for Alzheimer’s disease and autoimmune diseases. While laboratory model work has shown major longevity effects of single genes or single pathways, our results tend to support a model of human longevity influenced by multiple smaller effect protective and risk variants in several pathways, most prominently in cardiovascular related phenotypes. Our findings also supports the Geroscience concept that chronic diseases and biological processes involved in aging and longevity share similar mechanisms and pathways; efforts to target them may help both delay or prevent the onset of chronic diseases while also increasing longevity [23].

The *CHRNA3* rs1051730 variant has previously been linked to smoking fewer cigarettes and lower risks of lung cancer [21], although this variant does not influence the chances of starting smoking. Our finding of association with father’s age at death is highly plausible given the strongly adverse effect of smoking on survival. The fact that this SNP is also associated with mother’s age at death may support this finding, with the weaker (non-genome wide significant) association perhaps explained by the lower rates of smoking in women in the generation of parents being studied. We have noted that our attempt to test this variant independently in the Framingham Heart Study was not successful, but this analysis is severely underpowered, underlining the relatively small effect of this SNP on father’s longevity. Additionally, rs1051730 is located in a cluster of variants in high linkage disequilibrium covering several genes so the causal association could feasibly be via another mechanism (see **Supplementary Figure 6).**

We also analyzed associations with extreme longevity (here defined as at least one parent in the top 1% of age at death, ≥98 years for mothers or ≥95 years for fathers). The GWAS analysis revealed two suggestive genetic variants that require further study; rs528161076 is in a distinct peak of variants in an intron of *AP5Z1,* a gene encoding a homologous recombination DNA double-strand break repair helicase protein [22], and rs75824829 on chromosome 9 is ∼50k from the nearest gene *C9orf62 (chromosome 9 open reading frame 62)* for which little information is available. For the *AP5Z1* variant, this involves a C insertion after the T allele, with a prevalence 4.5% with at least one copy and is enriched amongst the those with top 1% longevity parents: of 1,335 participants with at least one long-lived parent, 84 have at least one of these insertions (6.3%) compared to 4.39% (1,796 of 40,871) of controls. Finding big enough independent samples to confirm these associations will be challenging.

For the genetic risk score analysis, several associations with continuous longevity become non-significant for extreme longevity, apparently due to lower sample sizes, but the pattern of GRS associations for both phenotypes were similar **(Figure 1).** More extreme definitions of longevity might yield different results but numbers of centenarian parents, for example, were too small to study. A possible exception to the generally similar risk pattern is the HDL cholesterol GRS, which was associated with extreme longevity but did not reach significance with continuous longevity. A number of other traits including type-1 diabetes GRS were close to significance, but these require additional study in a larger sample.

Associations between germline genetic variants and phenotypes provide strong evidence of causality because variants are inherited at the beginning of life, before any confounding by environmental or other factors can occur. Our results confirm the importance of several potentially manageable risk factors in achieving exceptional longevity, including adverse lipid levels, raised blood pressure (systolic and diastolic), adiposity (body mass index) and also smoking, as noted above. Although there has been some debate on whether there are paradoxical associations between obesity risks factors and survival in old age [24], our result suggests that genetically influenced life-long exposures in the expected directions (i.e. non-paradoxical effects) do influence human survival. This genetic evidence supports recent work on BMI and survival in a sample of 1 million older people, suggesting that paradoxical associations reflect reverse causation from weight loss in older people who already have major diseases associated with weight loss (Bowman et al, PLoS Medicine *in review* 2016).

### Comparison with previous work

There is consistent epidemiological evidence for a relationship between longevity and lower rates of type-2 diabetes [11], yet previous evidence from the Leiden study (n=2415) and the Life Long Family Study (n=1562 in generation 1; n=3102 generation 2) suggested that offspring of long-lived participants do not have lower burden of type-2 diabetes risk alleles compared to their partners [10,25]. We also did not observe an association between the type 2 diabetes GRS and longevity in the UK Biobank participants, although small effects cannot be excluded.

There is evidence to suggest that long-lived individuals have reduced prevalence of Alzheimer’s increasing alleles [10], which includes the *TOMM40/APOE* locus rs2075650 identified in multiple analyses of longevity [8,26,27]. We confirmed that the A allele of this SNP was positively associated with increased parental age at death (beta=0.039, p=1x10^-4^), however there was no significant association with “extreme longevity” (at least one parent in the top 1% of ages at death). The A allele, here associated with increased parents lifespan, is associated with decreased risk of dementia [28].

A recent study found that longevity-associated loci are enriched for disease-associated loci, consistent with our findings in Biobank [29]. The authors also performed a disease-weighted GWAS analysis, reporting 4 loci associated with exceptional longevity (≥90 years) with replication. We replicate the association between rs3184504 (mapped to *SH2B3/ATXN2),* previously shown to be associated with celiac disease [30], in our extreme longevity phenotype, but not the other three (although they are associated with continuous parent’s age at death p<0.05). rs3184504 has also been associated with blood pressure and cardiovascular disease [31]. This suggests rs3184504 is associated with survival to exceptional ages, but the others require further evidence.

Previous analyses have found that long-lived individuals have lower low-density lipoprotein genetic risk than young controls [32,33], which we also observe (offspring of long-lived parents have lower LDL-increasing genetic risk score). Our results extend this work by showing associations also with triglyceride and HDL genetic risk scores, plus BMI and systolic blood pressure, as well as associations with non-lipid cardiovascular disease genetic risk traits.

### Limitations

This study is limited to white British UK biobank participants of Caucasian genetic descent, thus the results may not be applicable to other populations. We plan to address this in future collaborations. In addition, the UK Biobank is a volunteer study that did not aim for population representativeness at baseline, although efforts were made to recruit a heterogeneous sample by varying geographic placement of examination sites, including in economically deprived areas; the final response rate was 5.47% [34]. It has been reported that with sufficient variation in the phenotypes being studied, results are generalizable to the wider UK population [35].

We are limited by the coverage of the genotyping microarray utilized; ∼800,000 genetic variants were directly measured, allowing imputation of 73 million (9 million of which were common enough and high quality enough for inclusion in this GWAS). Many variants may exist outside of the data available for this study, in particular on the X, Y and mitochondrial chromosomes, for which imputed data are not available.

The available information about the parents is very limited, with no dates of birth for those who had died and no cause of death information. We have excluded UK Biobank participants aged less than 55 to avoid adding large numbers of parents who are likely to be too young to have reached the longer lifespans which are our main focus in this analysis. The lack of cause of death data may have resulted in an underestimation of effect sizes on lifespan due to aging and related disease, as we have had to include e.g. accidental deaths unrelated to aging.

When comparing the effect sizes between different GRS, it is important to consider that each GRS explains a different proportion of its associated phenotype: this will in turn affect the interpretation of the association with longevity. For instance, only approximately 1% of the variance in mean length of the telomeres is explained by the 7 reported SNPs [36], whereas more than 10% of the variance in the blood lipids is accounted for by associated variants [37]; therefore the strength of association between these two phenotypes and longevity may not be proportionately represented by the effect sizes of the GRS longevity associations.

## Conclusions

Human longevity in our studied Caucasian origin sample is influenced by multiple common risk and protective genetic variants. Cardiovascular trait variants are particularly prominent in associations with longevity. Several of these genetically influenced risks including tobacco exposure are potentially modifiable. Further work is needed in other ethnic groups and to test less common variants for associations with longevity.

## Funding and Acknowledgements

This work was generously funded by an award to DM, TF, AM, LH and CB by the Medical Research Council MR/M023095/1. This research has been conducted using the UK Biobank Resource, under application 1417. The authors wish to thank the UK Biobank participants and coordinators for this unique dataset.

S.E.J. is funded by the Medical Research Council (grant: MR/M005070/1). J.T. is funded by a Diabetes Research and Wellness Foundation Fellowship. R.B. is funded by the Wellcome Trust and Royal Society grant: 104150/Z/14/Z. M.A.T., M.N.W. and A.M. are supported by the Wellcome Trust Institutional Strategic Support Award (WT097835MF). R.M.F. is a Sir Henry Dale Fellow (Wellcome Trust and Royal Society grant: 104150/Z/14/Z). A.R.W. H.Y., and T.M.F. are supported by the European Research Council grant: 323195:GLUCOSEGENES-FP7-IDEAS-ERC. The funders had no influence on study design, data collection and analysis, decision to publish, or preparation of the manuscript.

This work has made use of the resources provided by the University of Exeter Science Strategy and resulting Systems Biology initiative. Primarily these include high-performance computing facilities managed by Konrad Paszkiewicz of the College of Environmental and Life Sciences and Pete Leggett of the University of Exeter Academics services unit.

## References

[1] López-Otín C, Blasco MA, Partridge L, Serrano M, Kroemer G. The hallmarks of aging. Cell. 2013; 153:1194–217.

[2] Seok J, Warren HS, Cuenca AG, Mindrinos MN, Baker H V, Xu W, Richards DR, McDonald-Smith GP, Gao H, Hennessy L, Finnerty CC, López CM, Honari S, et al. Genomic responses in mouse models poorly mimic human inflammatory diseases. Proc Natl Acad Sci U S A. 2013; 110:3507–12.

[3] Melzer D, Frayling TM, Murray A, Hurst AJ, Harries LW, Song H, Khaw K, Luben R, Surtees PG, Bandinelli SS, Corsi A-M, Ferrucci L, Guralnik JM, et al. A common variant of the p16(INK4a) genetic region is associated with physical function in older people. Mech Ageing Dev. 2007; 128:370–7.

[4] Ostbye T, Taylor DH, Krause KM, Van Scoyoc L, Taylor Jr. DH, Krause KM, Van Scoyoc L. The Role of Smoking and Other Modifiable Lifestyle Risk Factors in Maintaining and Restoring Lower Body Mobility in Middle-Aged and Older Americans: Results from the HRS and AHEAD. J Am Geriatr Soc. 2002; 50:691.

[5] Herskind AM, McGue M, Holm N V, Sørensen TI, Harvald B, Vaupel JW. The heritability of human longevity: a population-based study of 2872 Danish twin pairs born 1870-1900. Hum Genet. 1996; 97:319–23.

[6] Mitchell BD, Hsueh WC, King TM, Pollin TI, Sorkin J, Agarwala R, Schäffer AA, Shuldiner AR. Heritability of life span in the Old Order Amish. Am J Med Genet. 2001; 102:346–52.

[7] Hjelmborg JB, Iachine I, Skytthe A, Vaupel JW, McGue M, Koskenvuo M, Kaprio J, Pedersen NL, Christensen K. Genetic influence on human lifespan and longevity. Hum Genet. 2006; 119:312–321.

[8] Broer L, Buchman AS, Deelen J, Evans DS, Faul JD, Lunetta KL, Sebastiani P, Smith J a, Smith A V, Tanaka T, Yu L, Arnold AM, Aspelund T, et al. GWAS of Longevity in CHARGE Consortium Confirms APOE and FOXO3 Candidacy. J Gerontol A Biol Sci Med Sci. 2014:1–9.

[9] Beekman M, Nederstigt C, Suchiman HED, Kremer D, van der Breggen R, Lakenberg N, Alemayehu WG, de Craen AJM, Westendorp RGJ, Boomsma DI, de Geus EJC, Houwing-Duistermaat JJ, Heijmans BT, et al. Genome-wide association study (GWAS)-identified disease risk alleles do not compromise human longevity. Proc Natl Acad Sci U S A. 2010; 107:18046–9.

[10] Stevenson M, Bae H, Schupf N, Andersen S, Zhang Q, Perls T, Sebastiani P. Burden of disease variants in participants of the Long Life Family Study. Aging (Albany NY). 2015; 7:123–32.

[11] Dutta A, Henley W, Robine JM, Langa KM, Wallace RB, Melzer D. Longer lived parents: Protective associations with cancer incidence and overall mortality. Journals Gerontol - Ser A Biol Sci Med Sci. 2013; 68:1409–1418.

[12] Tan Q, Zhao JH, Li S, Kruse TA, Christensen K. Power assessment for genetic association study of human longevity using offspring of long-lived subjects. Eur J Epidemiol. 2010; 25:501–6.

[13] UK Biobank. UK Biobank genotype collections and QC on 150k participants, April 2014; 2014. http://www.ukbiobank.ac.uk/wp-content/uploads/2014/04/UKBiobank_genotyping_QC_documentation-web.pdf.

[14] Abraham G, Inouye M. Fast principal component analysis of large-scale genome-wide data. PLoS One. 2014; 9:e93766.

[15] Gauderman W, Morrison J. QUANTO 1.1: A computer program for power and sample size calculations for genetic-epidemiology studies 2006.

[16] Loh P-R, Tucker G, Bulik-Sullivan BK, Vilhjálmsson BJ, Finucane HK, Salem RM, Chasman DI, Ridker PM, Neale BM, Berger B, Patterson N, Price AL. Efficient Bayesian mixed-model analysis increases association power in large cohorts. Nat Genet. 2015; 47:284–90.

[17] Purcell S, Neale B, Todd-Brown K, Thomas L, Ferreira MAR, Bender D, Maller J, Sklar P, de Bakker PIW, Daly MJ, Sham PC. PLINK: a tool set for whole-genome association and population-based linkage analyses. Am J Hum Genet. 2007; 81:559–75.

[18] Kannel WB, Feinleib M, McNamara PM, Garrison RJ, Castelli WP. An investigation of coronary heart disease in families. The Framingham offspring study. Am J Epidemiol. 1979; 110:281–90.

[19] Loh P-R, Bhatia G, Gusev A, Finucane HK, Bulik-Sullivan BK, Pollack SJ, Schizophrenia Working Group of the Psychiatric Genomics Consortium, de Candia TR, Lee SH, Wray NR, Kendler KS, O’Donovan MC, Neale BM, et al. Contrasting genetic architectures of schizophrenia and other complex diseases using fast variance-components analysis. Nat Genet. 2015; 47:1385–92.

[20] Pruim RJ, Welch RP, Sanna S, Teslovich TM, Chines PS, Gliedt TP, Boehnke M, Abecasis GR, Willer CJ. LocusZoom: regional visualization of genome-wide association scan results. Bioinformatics. 2010; 26:2336–7.

[21] Furberg H, Kim Y, Dackor J, Boerwinkle E, Franceschini N, Ardissino D, Bernardinelli L, Mannucci PM, Mauri F, Merlini PA, Absher D, Assimes TL, Fortmann SP, et al. Genome-wide meta-analyses identify multiple loci associated with smoking behavior. Nat Genet. 2010; 42:441–447.

[22] Rebhan M, Chalifa-Caspi V, Prilusky J, Lancet D. GeneCards: integrating information about genes, proteins and diseases. Trends Genet. 1997; 13:163.

[23] Kennedy BK, Berger SL, Brunet A, Campisi J, Cuervo AM, Epel ES, Franceschi C, Lithgow GJ, Morimoto RI, Pessin JE, Rando TA, Richardson A, Schadt EE, et al. Geroscience: Linking Aging to Chronic Disease. Cell. 2014; 159:709–713.

[24] Dixon JB, Egger GJ, Finkelstein EA, Kral JG, Lambert GW. ‘Obesity paradox’ misunderstands the biology of optimal weight throughout the life cycle. Int J Obes (Lond). 2015; 39:82–4.

[25] Mooijaart SP, van Heemst D, Noordam R, Rozing MP, Wijsman C a, de Craen AJM, Westendorp RGJ, Beekman M, Slagboom PE. Polymorphisms associated with type 2 diabetes in familial longevity: The Leiden Longevity Study. Aging (Albany, NY Online). 2011; 3:55–62.

[26] Deelen J, Beekman M, Uh H-W, Helmer Q, Kuningas M, Christiansen L, Kremer D, van der Breggen R, Suchiman HED, Lakenberg N, van den Akker EB, Passtoors WM, Tiemeier H, et al. Genome-wide association study identifies a single major locus contributing to survival into old age; the APOE locus revisited. Aging Cell. 2011; 10:686–98.

[27] Nebel A, Kleindorp R, Caliebe A, Nothnagel M, Blanché H, Junge O, Wittig M, Ellinghaus D, Flachsbart F, Wichmann H-E, Meitinger T, Nikolaus S, Franke A, et al. A genome-wide association study confirms APOE as the major gene influencing survival in long-lived individuals. Mech Ageing Dev. n.d.; 132:324–30.

[28] Yu C-E, Seltman H, Peskind ER, Galloway N, Zhou PX, Rosenthal E, Wijsman EM, Tsuang DW, Devlin B, Schellenberg GD. Comprehensive analysis of APOE and selected proximate markers for late-onset Alzheimer’s disease: patterns of linkage disequilibrium and disease/marker association. Genomics. 2007; 89:655–65.

[29] Fortney K, Dobriban E, Garagnani P, Pirazzini C, Monti D, Mari D, Atzmon G, Barzilai N, Franceschi C, Owen AB, Kim SK. Genome-Wide Scan Informed by Age-Related Disease Identifies Loci for Exceptional Human Longevity. PLOS Genet. 2015; 11:e1005728.

[30] Hunt KA, Zhernakova A, Turner G, Heap GAR, Franke L, Bruinenberg M, Romanos J, Dinesen LC, Ryan AW, Panesar D, Gwilliam R, Takeuchi F, McLaren WM, et al. Newly identified genetic risk variants for celiac disease related to the immune response. Nat Genet. 2008; 40:395–402.

[31] Huan. Integrative network analysis reveals molecular mechanisms of blood pressure regulation. Mol Syst Biol. 2015.

[32] Postmus I, Deelen J, Sedaghat S, Trompet S, de Craen a. J, Heijmans BT, Franco OH, Hofman a., Dehghan a., Slagboom PE, Westendorp RG, Jukema JW. LDL cholesterol still a problem in old age? A Mendelian randomization study. Int J Epidemiol. 2015; 73:604–612.

[33] Atzmon G, Rincon M, Schechter CB, Shuldiner AR, Lipton RB, Bergman A, Barzilai N. Lipoprotein Genotype and Conserved Pathway for Exceptional Longevity in Humans. PLoS Biol. 2006; 4:e113.

[34] Allen N, Sudlow C, Downey P, Peakman T, Danesh J, Elliott P, Gallacher J, Green J, Matthews P, Pell J, Sprosen T, Collins R. UK Biobank: Current status and what it means for epidemiology. Heal Policy Technol. 2015; 1:123–126.

[35] Manolio TA, Collins R. Enhancing the feasibility of large cohort studies. JAMA. 2010; 304:2290–1.

[36] Codd V, Nelson CP, Albrecht E, Mangino M, Deelen J, Buxton JL, Hottenga JJ, Fischer K, Esko T, Surakka I, Broer L, Nyholt DR, Leach IM, et al. Identification of seven loci affecting mean telomere length and their association with disease. Nat Genet. 2013; 45:422–427.

[37] Global Lipids Genetics Consortium, Willer CJ, Schmidt EM, Sengupta S, Peloso GM, Gustafsson S, Kanoni S, Ganna A, Chen J, Buchkovich ML, Mora S, Beckmann JS, Bragg-Gresham JL, et al. Discovery and refinement of loci associated with lipid levels. Nat Genet. 2013; 45:1274–83.

